# Estimation of substitution and indel rates via *k*-mer statistics

**DOI:** 10.1101/2025.05.14.653858

**Authors:** Mahmudur Rahman Hera, Paul Medvedev, David Koslicki, Antonio Blanca

## Abstract

Methods utilizing *k*-mers are widely used in bioinformatics, yet our understanding of their statistical properties under realistic mutation models remains incomplete. Previously, substitution-only mutation models have been considered to derive precise expectations and variances for mutated *k*-mers and intervals of mutated and non-mutated sequences. In this work, we consider a mutation model that incorporates insertions and deletions in addition to single-nucleotide substitutions. Within this framework, we derive closed-form *k*-mer-based estimators for the three fundamental mutation parameters: substitution, deletion rate, and insertion rates. We provide theoretical guarantees in the form of concentration inequalities, ensuring accuracy of our estimators under reasonable model assumptions. Empirical evaluations on simulated evolution of genomic sequences confirm our theoretical findings, demonstrating that accounting for insertions and deletions signals allows for accurate estimation of mutation rates and improves upon the results obtained by considering a substitution-only model. An implementation of estimating the mutation parameters from a pair of fasta files is available here: github.com/KoslickiLab/estimate_rates_using_mutation_model.git. The results presented in this manuscript can be reproduced using the code available here: github.com/KoslickiLab/est_rates_experiments.git.

**2012 ACM Subject Classification:** Applied computing → Computational biology; Theory of computation → Theory and algorithms for application domains; Mathematics of computing → Probabilistic inference problems

## 1 Introduction

Estimating the mutation rate between two evolutionarily related sequences is a classical question in molecular evolution, with roots that pre-date the genomics era [25]. Early quantitative efforts focused on amino-acid substitution: the seminal PAM matrices of Dayhoff et al. converted curated alignments of close homologues into an evolutionary time-scale [5], while the BLOSUM series by Henikoff and Henikoff mined ungapped blocks of conserved proteins to improve sensitivity for more diverged sequences [8]. These approaches and later profile-based HMM models [6] were derived from pairwise or multiple alignments and remain the gold standard when accurate alignments are available.

Over the last decade, however, high-throughput sequencing has shifted the scale of comparative genomics from dozens to millions of genomes, rendering high computational cost pipelines (e.g. quadratic-time) increasingly impractical. Consequently, alignment-free techniques that summarize sequences by inexpensive statistics have become indispensable [21, 26]. These approaches most commonly utilize *k*-mer sets and sketches thereof. Popular tools such as Mash [14], Skmer [17] and more recent sketch-corrected frameworks like Sylph [20] and FracMinHash-based methods [9, 11, 15, 19] can build whole-genome phylogenies, screen metagenomic samples, and compute millions of pairwise point-mutation rate estimates in minutes rather than days.

Despite their empirical success, theoretical understanding of alignment-free estimators has lagged behind practice. Nearly all existing models treat evolution as a pure-substitution process, ignoring insertions and deletions (indels), or else their performance in the presence of indels is often not thoroughly evaluated [16]. When indels are frequent, substitution-only estimators systematically inflate divergence and can misplace taxa—even in otherwise well-resolved trees of primates constructed from *k*-mer Jaccard similarities [14]. Recent work has quantified how *k*-mer-based statistics are affected by substitutions and are also used to estimate substitution-only mutation rates [11], yet a principled treatment that jointly infers substitution and indel parameters from *k*-mer statistics is still absent. This omission is particularly significant because indels represent a substantial fraction of genomic variation and play crucial roles in evolution [23]. Such indels cause substitution-only *k*-mer-based methods to underperform, as just like with substitutions, disruption of *k*-mer content by indel events affects at least *k k*-mers, often leading to overestimates of mutation rates [3, 11].

In this paper we introduce the first closed-form, alignment-free estimators for the three fundamental mutation parameters: substitution rate *p*_*s*_, deletion rate *p*_*d*_, and mean insertion length *g* under a model that explicitly incorporates single-nucleotide substitutions, deletions, and geometrically-distributed insertions. Starting from elementary counts of non-mutated and single-deletion *k*-mers, we derive algebraic expressions for *p*_*s*_, *p*_*d*_, and *g* and prove their accuracy via sub-exponential concentration bounds (Theorem 2). Simulations on synthetic and real bacterial genomes demonstrate that modeling indels yields markedly more accurate distance estimates than substitution-only approaches. The remainder of this paper is organized as follows: Section 2 gives preliminary definitions and defines our mutation model, Section 3 presents our main results, including the estimators and their theoretical guarantees, Section 4 provides a proof outline of the main theorem, Section 5 details our implementation, Section 6 presents experimental results, and Section 7 concludes with a discussion of implications and future directions. The Appendix contains most of the proofs.

## 2 Preliminaries

We will use *S* to denote a string over the alphabet {*A, C, G, T* }. We define *L* to denote the number of characters in *S*. In this paper, we assume that there is a fixed integer 1 ≤ *k* ≤ *L*, which forms the basis of our analysis. We will use *L*_0_ as a shorthand for *L* − *k* + 1. We will also assume that *L* ≥ *k*. We use *S* _*i*_ to denote the *i*-th character in *S*, where 1 ≤ *i* ≤ *L*. Let *c*_*A*_ be the number of ‘A’s in *S*.

We use the mutation model from [18]. This model is a type of indel channel and various variations of it have been used to model sequence evolution (e.g. [7]). The mutation process takes as input a string *S* and three parameters *p*_*s*_, *p*_*d*_, and *g*, where 0 ≤ *p*_*s*_, *p*_*d*_ < 1, *p*_*s*_ + *p*_*d*_ < 1, and *g* ≥ 0. It performs the following steps on *S*:

1. For each *i*, independently choose an operation from the set {Sub, Del, Stay} with respective probabilities *p*_*s*_, *p*_*d*_, and 1 − *p*_*s*_ − *p*_*d*_.
2. Let *track* be a function mapping from a position in the original string *S* to its position in the modified *S*. Initially, *track*(*i*) = *i* for all *i*. We assume that *track* is updated accordingly whenever *S* will be modified by either deleting or inserting characters.
3. For each *i* with a Sub operation, select a character *c* ∈ {*A, C, G, T* } \ {*S* _*i*_} uniformly at random and replace *S* _*i*_ by *c*.
4. For each *i*, let *I*_*i*_ ≥ 0 be a sample from a geometric distribution with mean *g*. Then generate a random string of length exactly *I*_*i*_ by drawing each character of the string from {*A, C, G, T* } independently and uniformly at random. We call this the *insert string associated with position i*. If *I*_*i*_ = 0, then this is the empty string.
5. For every *i* > 1, insert the string associated with position *i* between *S* _*track*(*i*)−1_ and *S* _*track*(*i*)_, updating *track* accordingly after each insertion.
6. Prepend the insert string associated with position 1 to the start of *S*, updating *track* accordingly.
7. For every *i* with a Del operation, remove the character *S* _*track*(*i*)_ from *S*, updating *track* accordingly after each deletion.

A *k*-span is an interval of length *k* on the interval [1, *L*]. We define *K*_*i*_ as the *k*-span starting at position 1 ≤ *i* ≤ *L*_0_, i.e. the interval [*i, i* + *k* − 1]. We say that *K*_*i*_ *contains a mutation* if there exists a *i* ≤ *j* < *i* + *k* with an operation other than Stay or there exists a *i* < *j* < *i* + *k* such that *l*_*i*_ * 0.

We use *S*′ to denote the random string output after the mutation process is applied on *S*, and we define 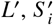, and *c*_*A*_′ for *S*′ analogously to how they are defined for *S*. We will also make use of three random variables based on the mutation process.

▄ *N* ≜ the number of *k*-spans that have no mutation.
▄ *D* ≜ the number of *k*-spans that have no mutation except for exactly one deletion.
▄ *P* ≜ *L*′ − 4*c*_*A*_′

## 3 Main result

For the remainder of the manuscript, we assume that the mutation process described in Section 2 is applied to string *S* of length *L*, with the unknown parameters *p*_*s*_, *p*_*d*_, and *g*, and a mutated string *S*′ is returned. Rather than observing the full string *S*′, we limit ourselves to deriving estimators from *L*′, *c*_*A*_′, *D*, and *N*. We assume that *L*′ and *c*_*A*_′ can be directly observed from the data, but we postpone the discussion on how we can observe *D* and *N* until Section 5.

To derive our estimators, we take the following approach. We derive the expected values of the observed variables as a function of our model parameters, and then plug in the observed variables in place of their expectations and solve for the model parameters. The expectations of the observed variables are straightforward to derive and given by the following lemma.

### ►Lemma 1.

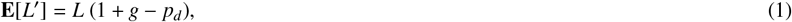

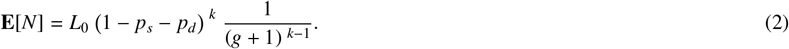

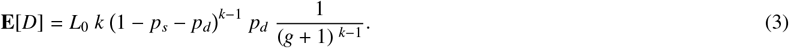

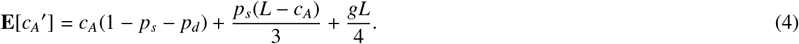

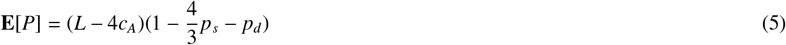

Next, we algebraically manipulate the equations in Lemma 1 in order to get formulas for *p*_*s*_, *p*_*d*_, and *g*. From Lemma 1 we get

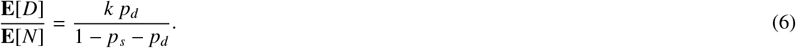

Note that we are not dividing by zero because our mutation model assumes *p*_*s*_ + *p*_*d*_ < 1. From (1), (4) and (6), we obtain the following system of linear equations with variables *p*_*s*_, *p*_*d*_, and *g*.

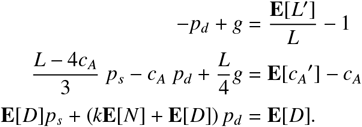

Solving this system of equations, we obtain that:

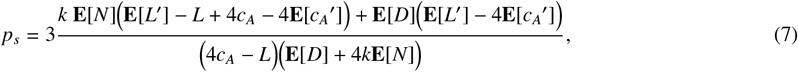

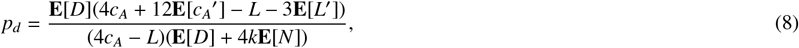

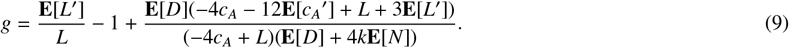

By replacing the expectations above with these observations, we obtain our estimators. That is,

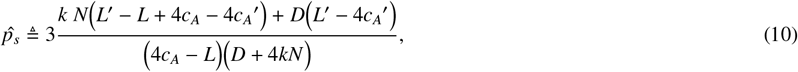

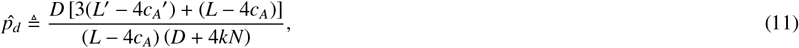

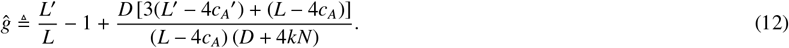

We briefly comment on our choice of estimators, as various statistical approaches based on a different set of observables could yield a different set of estimators. For example, we considered a variant of the estimators based on the counts of *k*-spans with a single insertion, substitution, or deletion (i.e. *D*). These quantities contain enough information to estimate the mutation parameters, specifically, by solving the non-linear system of equations given by (3) and similarly derived formulas for the expected number of *k*-spans with a single insertion and a single substitution; the resulting estimators performed quite well in real data. However, establishing theoretical guarantees for these estimators proved challenging, as they were defined as roots of degree-*k* polynomials. Our current estimators address this theoretical limitation as they involve solely linear equations. As we shall see in Section 6, the performance of our estimators in real data is strong, and they strike a more favorable balance by offering both reasonable accuracy and rigorous theoretical guarantees.

We now provide our central theoretical result. Our main theorem, below, shows that our estimates of *p*_*s*_, *p*_*d*_, and *g* are concentrated in a symmetric interval around their true values.

### ►Theorem 2.

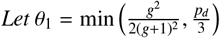 *and θ*_2_ = max(*g* + 1, 8). *Suppose* 4*c*_*A*_ < *L and* 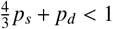. *Then, for δ*∈ (0, 1/5):

1. 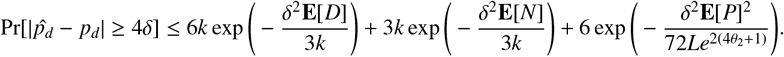
2. 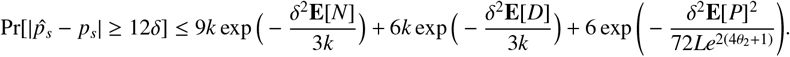
3. 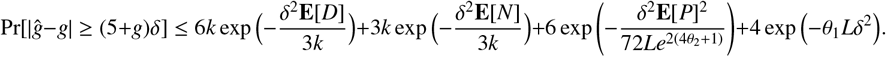

The requirement that 4*c*_*A*_ < *L* in this theorem does not restrict generality: aside from equal nucleotide frequency (where the estimators are not defined), at least one character *c* ∈ {*A, C, T, G*} must satisfy that 4 times its frequency is at most *L*. In addition, the assumptions that 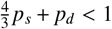 and *p*_*d*_ ≤ 1/2 hold when *p*_*s*_ and *p*_*d*_ are small which is the case most frequently encountered in practice. They are also non-essential and are taken to simplify our theoretical proofs.

The error probability in Theorem 2 is small when each of the terms in the sum are small. Since each of these terms decays (at least) exponentially with *δ*^2^ times an expectation that grows linearly with the length of the string (when the mutation parameters are fixed), they will all generally be small. The following corollary of Theorem 2 formalizes this idea.

### ►Corollary 3.

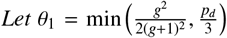 *and θ*_2_= max(*g* + 1, 8). *Suppose there exists ε* ∈ (0, 1) *independent of L such that* (4 + ε)*c*_*A*_ < *L, and* 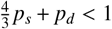. *Suppose further that p*_*s*_, *p*_*d*_, *g, and k are constants independent of L. Then, there exist constants γ* = *γ*(*p*_*s*_, *p*_*d*_, *g*, ε, *k*) > 0 *and C* = *C*(*k*) > 0 *such that for any δ* ∈ (0, 1/5):

1. 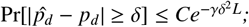
2. 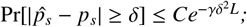
3. 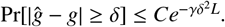

## 4 Proof outline of main theorem and corollary

In this section, we will outline the proof of the main theorem, with the missing proofs presented in Appendix B. We start by rewriting (7)-(12) in a more convenient form. Recall that 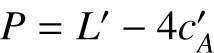 and define the following additional three variables:

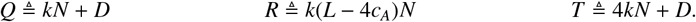

Applying these definitions to (7)-(12), we get the following alternate formulations for the true and estimated parameters.

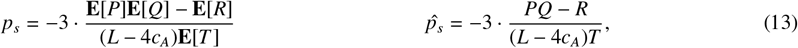

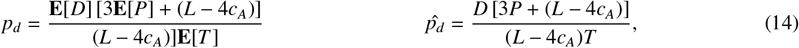

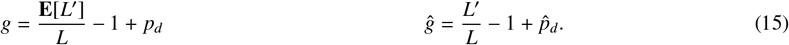

In order to derive the concentration bounds of Theorem 2, we will first derive concentration bounds for *P, Q, R, T*, and *L*′. *Q, R*, and *T* are all linear combinations of *N* and *D* and their concentration bounds follow easily once the concentration bounds for *N* and *D* are derived. The concentration bounds for *N* and *D* stem from the fact that, from the perspective of the mutation process *k*-spans located more than *k* positions apart are independent of each other. We can therefore partition the set of *L*_0_ *k*-spans into *k* subsets, such that the *k*-spans in each subset are independent of each other. We can then apply Chernoff bounds to each part and combine these bounds together via a union bound into a bound for *N*. In this way, we can obtain the following lemma.

### ►Lemma 4.

*Suppose that* 4*c*_*A*_ < *L. For any δ* ∈ (0, 1), *all the following hold:*

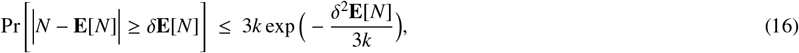

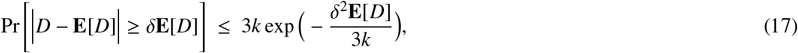

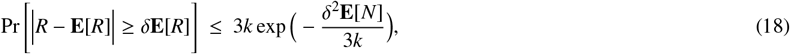

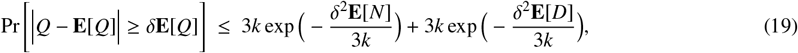

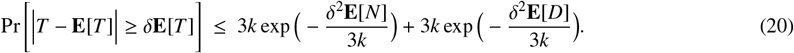

To prove concentration for the length of the mutated string *L*′, we separately bound the number of total inserted and deleted characters, which are negative binomially (i.e. sum of geometrically distributed variables) and binomially distributed, respectively.

### ►Lemma 5.

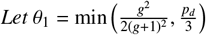. *For any δ* ∈ (0, 1):

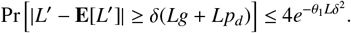

The proof of the concentration bound for *P* is more challenging, as *P* is a sum of independent but unbounded random variables and therefore the Chernoff and Hoeffding bounds cannot be applied directly. However, we are able to show that *P* can be expressed as the sum of independent random variables that are sub-exponential. The concentration of such sums can be bounded via Bernstein inequalities. We first derive a clean form of Bernstein’s inequality well-suited to our setting.

### ►Lemma 6.

*Let θ*_2_ = max(*g* + 1, 8). *Then, the following holds for any δ* > 0:

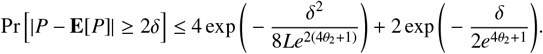

*If we also assume that* 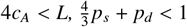, *and for any δ*_0_ ∈ (0, 1) *then*

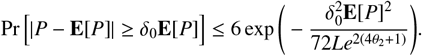

In order to combine these individual concentration inequalities into concentration inequalities for our estimators, which are ratios, we make use of the following technical lemma.

### ►Lemma 7.

*Let X, Y, and Z be random variables with strictly positive means. Let δ* ∈ (0, 1/5). *Suppose that* (1 − *δ*) **E**[*X*] ≤ *X* ≤ (1 + *δ*) **E**[*X*] *and* (1 − *δ*) **E**[*Y*] ≤ *Y* ≤ (1 + *δ*) **E**[*Y*]. *Then*

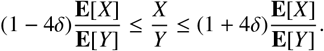

*If it also holds that* (1 − *δ*) **E**[*Z*] ≤ *Z* ≤ (1 + *δ*) **E**[*Z*], *then*

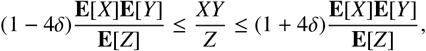

We can now prove Theorem 2. Here, we give the proof of the concentration of 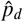, and leave the rest for Appendix B.

**Proof of Theorem 2, for** 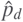. Consider the case that *D, P*, and *T* are indeed close to their means, i.e. for all *X* ∈ {*D, P, T* } and all *δ* ∈ (0, 1/5), we have (1 −*δ*) **E**[*X*] ≤ *X* ≤ (1 +*δ*) **E**[*X*]. Because we assume that 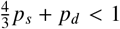 and 4*c*_*A*_ < *L*, we use (5) to get that **E**[*P*] > 0; Lemma 1 gives **E**[*D*], **E**[*T*] > 0. Therefore, we can apply Lemma 7 to the definition of 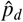 (14) to get

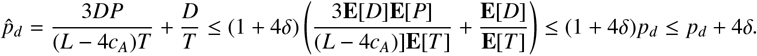

In a symmetric manner, we obtain that 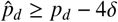. Hence, under the assumptions that *P, D*, and *T*, are close to their means, 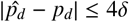. Conversely, when 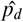 is outside this range, at least one of these three variables must be far from the mean. That is,

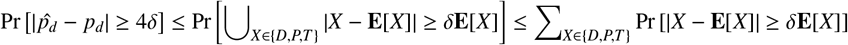

by a union bound. The result for 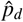 follows by plugging in the individual concentration bounds for *P* (Lemma 6), *Q*, and *T* (Lemma 4). ◀

**Proof of Corollary 3**. We prove the result for 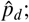 the results for 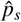 and *ĝ* follow analogously. Under the assumption that the model parameters are fixed and independent of *L*, it follows from Lemma 1 that there exist positive constants *γ*_*D*_ = *γ*_*D*_(*p*_*s*_, *p*_*d*_, *g, k*), *γ*_*N*_ = *γ*_*N*_ (*p*_*s*_, *p*_*d*_, *g, k*), and *γ*_*P*_ = *γ*_*P*_(*p*_*s*_, *p*_*d*_, ε) independent of *L* such that **E**[*D*] ≥ *γ*_*D*_*L*, **E**[*N*] ≥ *γ*_*N*_ *L*, and **E**[*P*] ≥ *γ*_*P*_*L*. Plugging these bounds into part 1 of Theorem 2, we obtain:

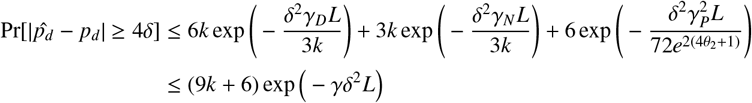

where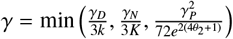. ◀

## 5 Implementation details

Our estimators for the three rates require counting the number of *k*-spans with single deletion, *D*, and the number of *k*-spans with no mutation, *N*. Counting *D* and *N* can be challenging, particularly because *k*-mers do not contain the contextual information, and so we do not have access to their corresponding *k*-spans. An additional layer of complexity comes into play from the fact that identifying a *k*-mer with no mutation (or a single kind of mutation) is more difficult, considering many edge cases that may arise from inserting the same character that has been deleted. These challenges are cimcumvented when we use *k*-spans, and therefore, counting *D* and *N* solely from *k*-mers is not trivial and can be considered an interesting problem in and of itself. We therefore implemented an ad hoc solution to estimate *D* and *N* given the two strings *S* and *S*′. The steps for estimating *D* and *N* are as follows.

We start by extracting all *k*-mers in *S*, and building a de Bruijn graph using these *k*-mers using the cuttlefish tool [13]. We then extract the unitigs from this graph. Let the set of unitigs computed from *S* be *U*. We also compute the unitigs in *S*′ in a similar manner and call this *U*′. We next take an arbitrary unitig *u* from *U*, and align every unitig in *U*′ with *u*. To allow for partial overlap, we use semi-global alignment by using the infix option in edlib [22], which makes sure that gaps at the beginning and at the end of the alignment are not penalized. For a particular *u* ∈ *U*, we align every *u*′ ∈ *U*′ to make sure all relevant alignments are considered. We use these alignments to look at all windows of length *k*, and count *D* and *N* accordingly for *u*. We repeat this for all *u* ∈ *U*, and accumulate the measurements from individual *u*’s into a single global count.

The motivation behind using unitigs is that if there is an isolated mutation, and if the mutation is in the first or the last position of a *k*-mer, then there is no way to understand if the mutation is a substitution, an insertion, or a deletion only from the *k*-mers. The only way to resolve this ambiguity (and other similar ambiguities) is to scan beyond the context of *k* characters – and unitigs are a natural way to do this. The core goal of our implementation of these steps described above was not to make it efficient, but rather to obtain a working solution. We found that executing these steps estimates *D* and *N* reasonably well, and the estimated rates are also acceptable. As such, we leave finding an efficient way to compute *D* and *N* as an open research question.

## 6 Experiments and results

In this section, we present a series of experiments to evaluate the performance of the estimators detailed in Section 3. As discussed earlier, these estimators are sensitive to several input parameters, including *k*-mer size, sequence length, and the fraction of ‘A’ characters in the sequence. Sections 6.1 through 6.3 explore the sensitivity of the estimators with respect to these parameters. In Sections 6.4 and 6.5, we estimate mutation rates across a wide range of known rate combinations. And finally, in Section 6.6, we demonstrate that our estimated substitution rate outperforms estimates obtained under a substitution-only mutation model. For the experiments in Sections 6.1 through 6.4, the original sequence is a randomly generated synthetic sequence. In these cases, we compute the number of *k*-spans containing a single deletion and the number of *k*-spans with no mutation directly from the known mutation process. For the experiments in Sections 6.5 and 6.6, we use real reference genomes as the original sequences. In these cases, the two types of *k*-span counts are estimated using the steps described in Section 5.

### 6.1 Sensitivity of the estimators to *k*-mer lengths

We begin our analysis by examining how the choice of *k*-mer length affects the estimation of the mutation rates. To investigate this effect, we first generated a synthetic reference sequence of 1 million nucleotides, randomly sampling bases with fixed frequencies: 30% ‘A’, and equal proportions of ‘C’, ‘G’, and ‘T’ – making sure total frequency is 100%. From this reference, we simulated 20 mutated sequences, independently from each other, using the mutation model described in Section 2. For each of these mutated sequences, we estimated mutation rates using the estimators defined in Section 3 using a range of values for *k*. The results of this analysis are summarized in Figure 1.

**Figure 1.**
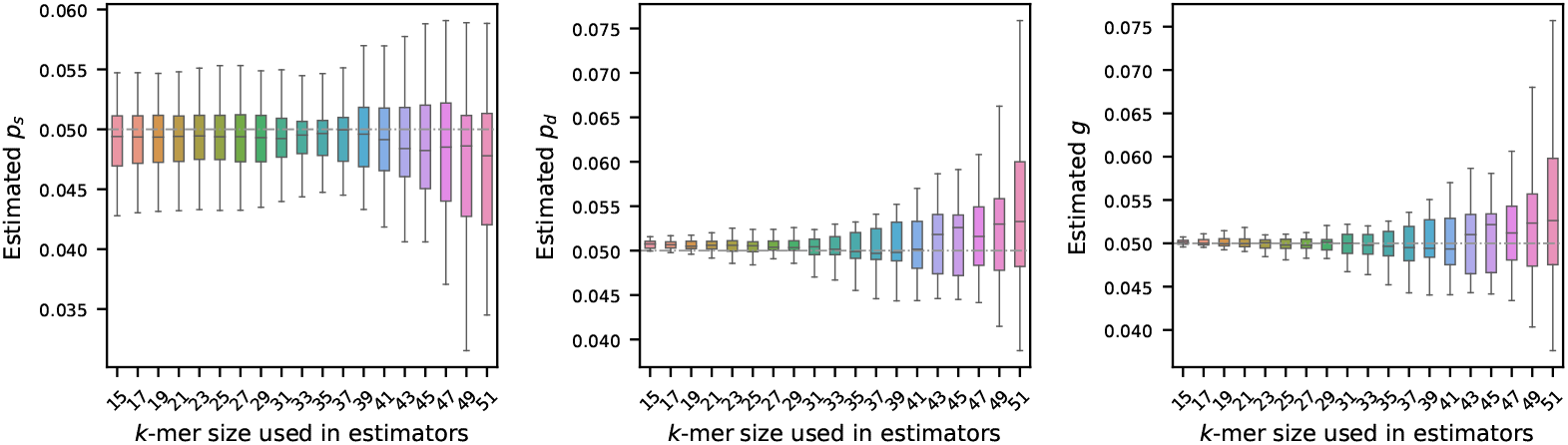
Effect of k-mer length on mutation rate estimation (true rates set to 0.05). A synthetic genome of 1 million nucleotides, mutated genomes were generated by setting *p*_*s*_, *p*_*d*_, and *g* = 0.05 – shown by the gray dashed horizontal line. Estimated rates were then computed using a range of *k*-mer sizes. Each boxplot shows the variability in estimation across 20 simulations, with error bars showing one standard deviation. The plots show that the estimators become more accurate and more precise for shorter *k*-mers.

As illustrated, the choice of *k* has a substantial impact on the stability of the estimators. In particular, longer *k*-mers tend to produce estimates with higher variability. This behavior is consistent with the known sensitivity of *k*-mers to mutations: since a single mutation can disrupt up to *k* consecutive *k*-mers, the longer the *k*-mer, the more susceptible it becomes to such perturbations. Our theoretical result in Theorem 2 also captures this: with a larger *k*, the error probabilities become larger, and the probabilistic guarantee for the estimators’ performances decreases accordingly.

Interestingly, for the estimator of substitution probability *p*_*s*_, we observe that the variability in the estimated values does not change significantly from 15 to 39. The reason behind this behavior is not immediately clear and warrants further investigation. It is possible that incorporating the number of *k*-spans with a single substitution into the estimators may correct this behavior, but additional analyses are required to substantiate this hypothesis.

### 6.2 Sensitivity of the estimators to sequence length

To investigate how the length of the original sequence *S* influences estimation of the mutation rates, we simulated synthetic genomes ranging from 10K to 1M nucleotides in length. For each genome length, we generated 10 independent synthetic sequences to capture variability due to random sampling. The nucleotide composition of each sequence was fixed, with the frequency of ‘A’ set to 30% and frequencies of ‘C’, ‘G’, and ‘T’ set equally – making sure total frequency is 100%. For each synthetic sequence, we generated its mutated version by running the mutation process described in Section 2, setting each of *p*_*s*_, *p*_*d*_, and *g* to 0.05. We then estimated the mutation rates using the estimators outlined in Section 3 for three *k*-mer sizes: *k* = 21, 31, 41.

Figure 2 displays the estimated rates across the varying sequence lengths. As shown, the estimators are less stable for shorter sequences. However, with longer sequences, the estimators yield more accurate results – a trend expected from our core theoretical result in Theorem 2, which states that the associated error is asymptotically vanishing in *L*, the length of the string *S*: as *L* increases, the number of *k*-spans increases, and therefore the probability of error decreases, leading to a more precise estimation.

**Figure 2.**
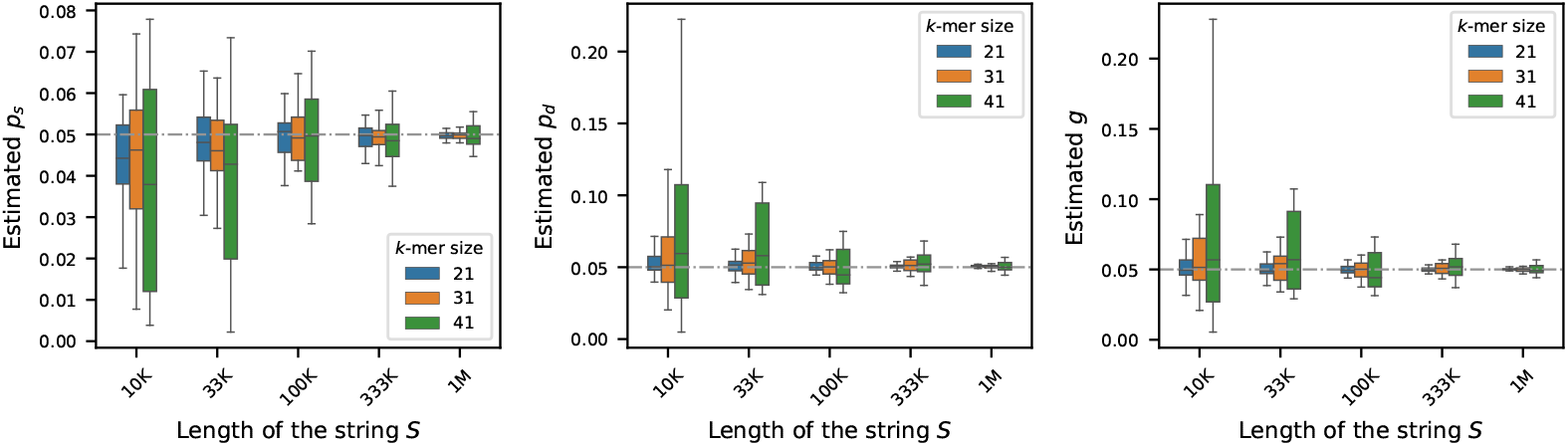
Effect of sequence length on mutation rate estimation (true rates set to 0.05). For synthetic genomes of varying lengths, mutated genomes were generated by setting *p*_*s*_, *p*_*d*_, and *g* = 0.05 – shown by the gray dashed horizontal line. Estimated rates were computed using three *k*-mer sizes: 21, 31, and 41. Each boxplot shows the variability in estimation across 20 simulations. The plots show that the estimators become more accurate and more precise for longer sequences.

### 6.3 Sensitivity of the estimators to base composition

We generated synthetic genomes of 1M nucleotides to investigate how the fraction of ‘A’ characters affects the estimation of the mutation rates. We varied the fraction of ‘A’s from 21% to 29% in increments of 1%. For each fraction of ‘A’s, we set the frequency of ‘C’s, ‘G’s, and ‘T’s equally. For each preset fraction of ‘A’s, we generated 10 random genomes to capture stochastic variation. For each of these genomes, we generated its mutated version using the mutation process described in Section 2, setting each of *p*_*s*_, *p*_*d*_, and *g* to 0.05. We then estimated the mutation rates using the estimators described in Section 3 using three *k*-mer sizes: *k* = 21, 31, and 41. Figure 3 shows the sensitivity of the estimators to the fraction of ‘A’s in the original string *S*.

**Figure 3.**
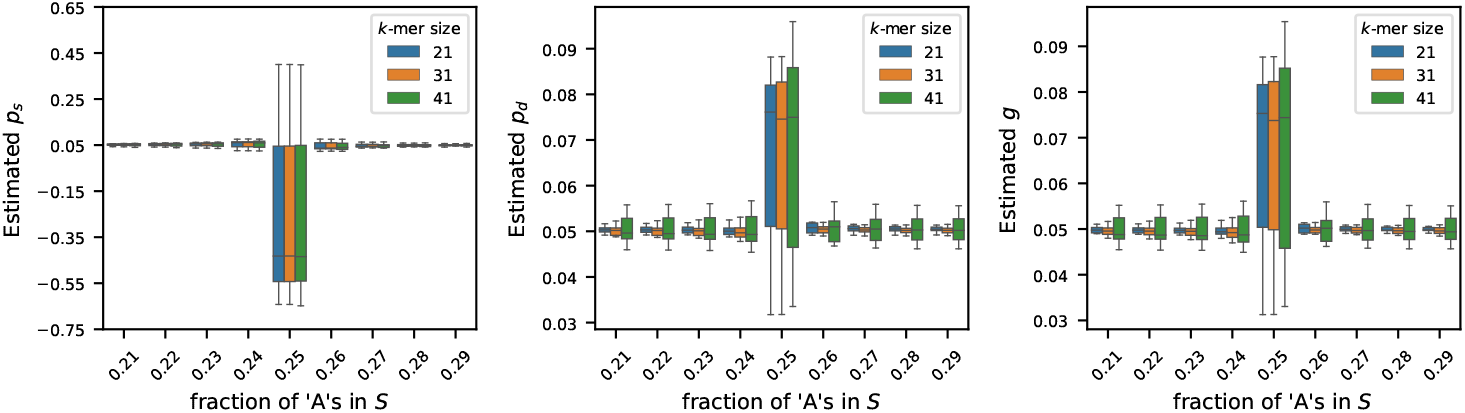
Effect of nucleotide composition on mutation rate estimation (true rates set to 0.05). For synthetic genomes of length *L* = 1M, the fraction of ‘A’s is varied from 0.21 to 0.29, and the frequencies of ‘G’s, ‘C’s, and ‘T’s are set equally. For each setting, the mutated string *S*′ was generated by setting *p*_*s*_, *p*_*d*_, *g* = 0.05. Estimated rates were computed for three *k*-mer sizes: 21, 31, and 41. Each boxplot shows the variability in estimation across 20 simulations. The results show that the estimators generally work well for all three *k*-mer sizes, except when *c*_*A*_ ≈ *L*/4, in which case the estimators become unstable – as predicted by Theorem 2.

We observe that the estimators 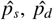, and *ĝ* work reasonably well to estimate the true rates when the frequency of ‘A’ characters, *c*_*A*_ is not *L*/4. On the contrary, when the fraction of ‘A’s is exactly 25% in the original string *S*, the estimator gives inaccurate values, some of which are even negative (see estimated values of *p*_*s*_). This behavior is captured in Theorem 2: when *L* − 4*c*_*A*_ = 0,*µ*_*P*_ = **E**[*P*] = 0, and the probabilistic guarantees become unbounded. We only get meaningful guarantees of concentration when 4*c*_*A*_ is strictly smaller than *L*. While Theorem 2 does not guarantee concentration when 4*c*_*A*_ > *L*, this does not restrict generality, as explained in Section 3, and as demonstrated by the estimators’ performances when the fraction of ‘A’s is larger than 1/4.

### 6.4 Estimating rates from a randomly generated synthetic sequence

After testing our estimators for varying *k*-mer lengths, sequence lengths, and base compositions, we next turn to estimating mutation rates by varying the true rates across a range of values. To do this, we generated a synthetic reference genome of 1 million base pairs, fixing the base composition at 30% ‘A’, and equal proportions of ‘C’, ‘G’, and ‘T’ – making sure total frequency is 100%. Using the mutation model described in Section 2, we then simulated mutated genomes from the synthetic reference by varying the mutation rates *p*_*s*_, *p*_*d*_, and *g* across the values {0.01, 0.02, 0.03, 0.04, 0.05}. For every parameter combination, we generated 10 independent mutated genomes to capture stochastic variability. We then estimated the mutation rates using the estimators detailed in Section 3 for each of these mutated genomes.

In Figure 4, we show two sets of results:

**Figure 4.**
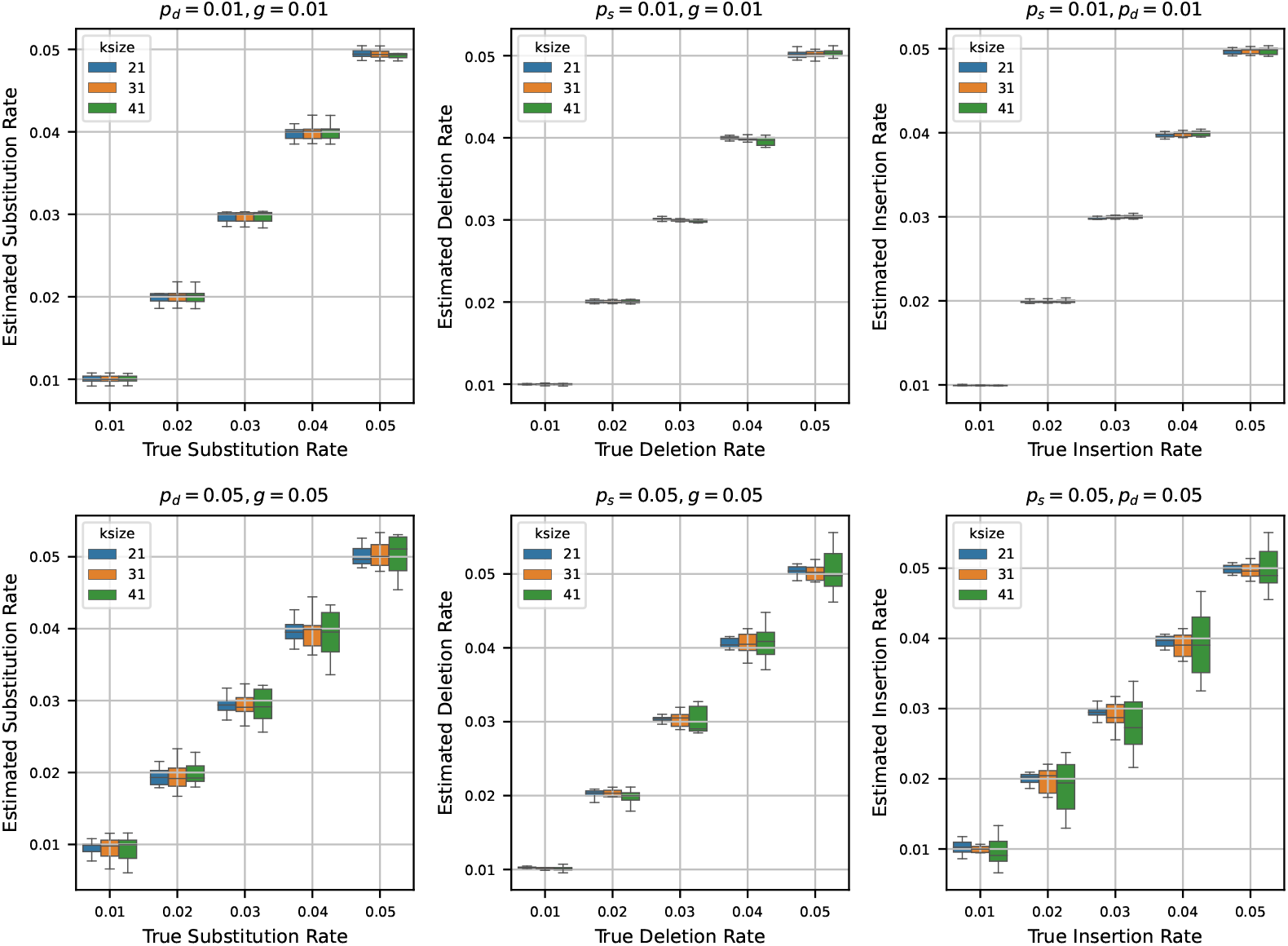
Estimated mutation rates versus true values, where the original string is a synthetic sequence. Each subplot corresponds to a case where two mutation rates are fixed (either at 0.01 or 0.05) and the third is varied from 0.01 to 0.05. Each boxplot shows the variability in estimation across 10 simulations. The results show that the rate estimation is very accurate when the other two rates are small, and is reasonably accurate when the other two rates are larger.

▄ **Fixed low rates (0.01):** Fixing two of the rates at 0.01, we show the estimates of the third rate as the true rate varies from 0.01 to 0.05. We repeat this process independently for *p*_*s*_, *p*_*d*_, and *g*.
▄ **Fixed high rates (0.05):** Fixing two of the rates at a higher value of 0.05, we also show the estimates of the third rate as the true rate varies from 0.01 to 0.05. Again, we repeat this for all three rates.

When we set the other two rates to 0.01 and estimate the third rate, we observe that the estimated rates are highly accurate across all trials. In many cases, the boxplots of the estimates nearly vanish due to minimal variance, indicating tight clustering around the true values. This trend remains consistent across multiple *k*-mer sizes, suggesting that the estimators are robust at low rates of mutation.

In contrast, when we fix the other two rates at 0.05 and estimate the third rate, the accuracy of the estimation decreases slightly. While the estimates still track the true values reasonably well, the variance increases, and the boxplots become more prominent. Notably, the median estimate remains close to the true rate in most settings, which indicates that the estimators retain their central tendency even under higher mutation rates. However, for larger *k*-mer sizes, we observe increased variability in the estimates – an effect that mirrors our earlier observations in Section 6.1, where longer *k*-mers resulted in decreased precision of the estimators.

### 6.5 Estimating rates from real sequences

Having the estimators tested for a synthetic reference, we next estimate rates from a real genome sequence. For this set of experiments, we used the reference assembly of *Staphylococcus aureus* (subspecies: aureus USA300_TCH1516), which has 2.8 million nucleotides. We simulated mutated sequences from this reference by running the mutation process described in Section 2 by varying the mutation rates *p*_*s*_, *p*_*d*_, and *g* from 0.01 to 0.05. Similar to Section 6.4, we generated 10 independent mutated sequences for each combination of *p*_*s*_, *p*_*d*_, and *g* to capture stochastic variability. We then estimated the mutation rates using the estimators outlined in Section 3 for each of these simulated sequences.

Figure 5 shows the estimated mutation rates plotted against the true rates that were used to run the mutation process. We observe that the results shown in Figure 5 are consistent with previously discussed results. Specifically, when estimating a given mutation rate while keeping the other two rates low (0.01), the corresponding estimator performs with high precision, closely tracking the true value. On the other hand, when the estimation is carried out with the other two mutation rates set to higher values (0.05), the estimates appear more confounded. This is likely due to the increased difficulty of accurately estimating the number of *k*-spans with a single deletion (*D*) or no mutation (*N*) in a real genomic context. Notably, these experiments involve challenging conditions, with total mutation rates exceeding 10%. Despite this, the estimators yield reasonably accurate results, indicating potential for practical effectiveness.

**Figure 5.**
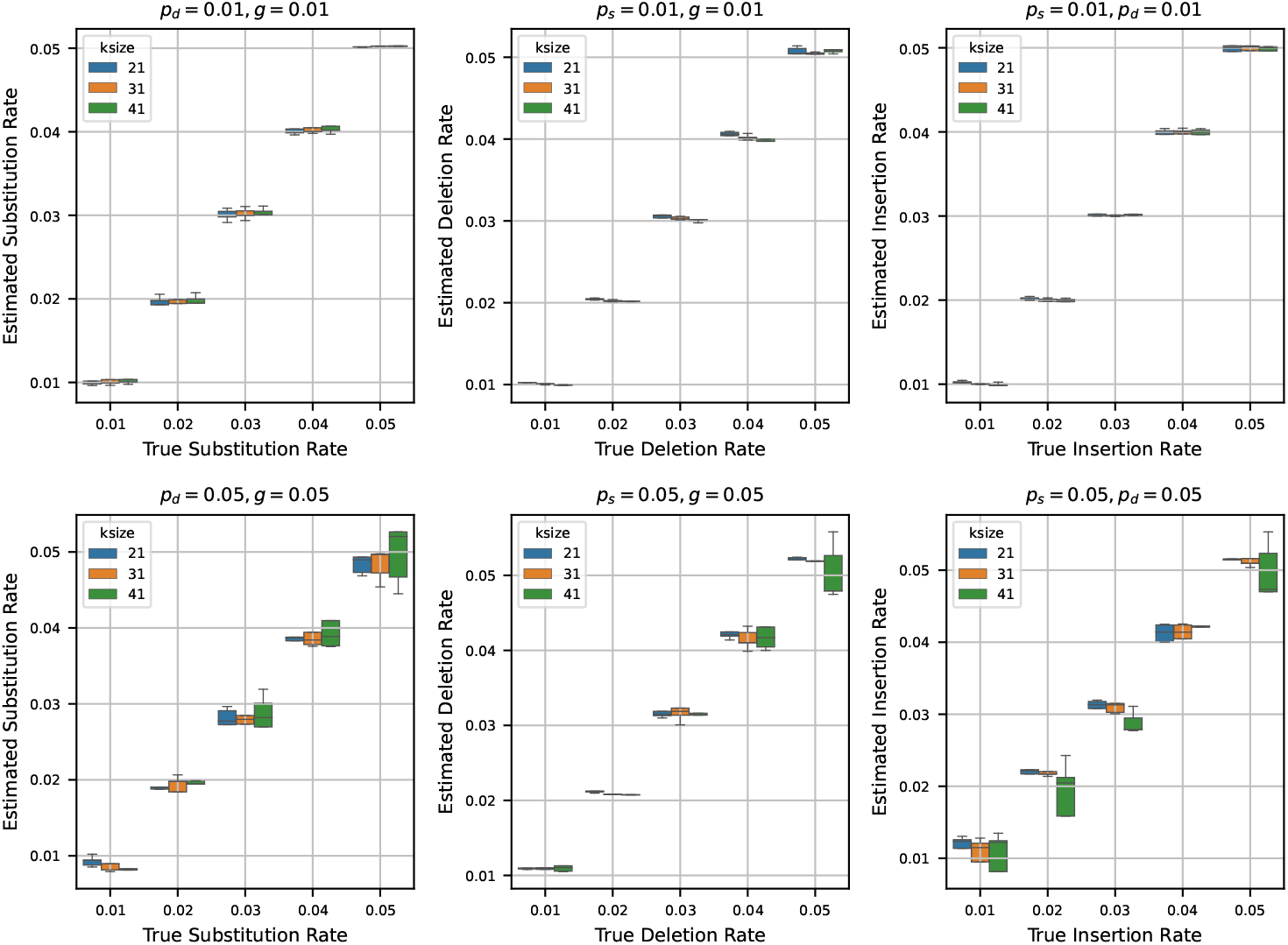
Estimated mutation rates versus true values, where the original string is the reference genome of Staphylococcus (length 2.8 million). Each subplot corresponds to a case where two mutation rates are fixed (either at 0.01 or 0.05) and the third varies from 0.01 to 0.05. Each boxplot shows the variability in estimation across 10 simulations. Estimated mutation rates closely match true values when other rates are low, but estimation becomes less precise under high total mutation rates (>10%) due to increased difficulty in real genomic contexts.

### 6.6 Comparison with substitution rates estimated using simple mutation model

We conclude the experiments section by contrasting the substitution rates estimated using (10) with substitution rates estimated considering a simple mutation model. We use the statistics of *k*-mers developed in a recent work [2] to estimate substitution rates under a simple mutation model. The simple mutation model captures only substitutions, and no insertions or deletions. Consequently, we can only compute substitution rates considering this simple model. Henceforth, we refer to the simple mutation model as SMM.

We estimated the substitution rates using the SMM for the same simulated mutated sequences described in Section 6.5. In Figure 6, we show the estimated substitution rates using our estimators in (10), and using SMM. The results highlight that the substitution rates estimated using the estimator we developed track the true substitution rates accurately. On the other hand, the substitution rates estimated using SMM make a gross overestimation. This is because the SMM does not consider indels, and therefore, the effects of all three mutation rates are subsumed in the single substitution rate we get using the SMM. As such, a simple mutation model cannot disentangle the distinct contributions of substitution, insertion, and deletion rates. In contrast, the mutation model we introduce effectively decomposes these components, enabling more accurate and meaningful estimation of individual mutation rates.

**Figure 6.**
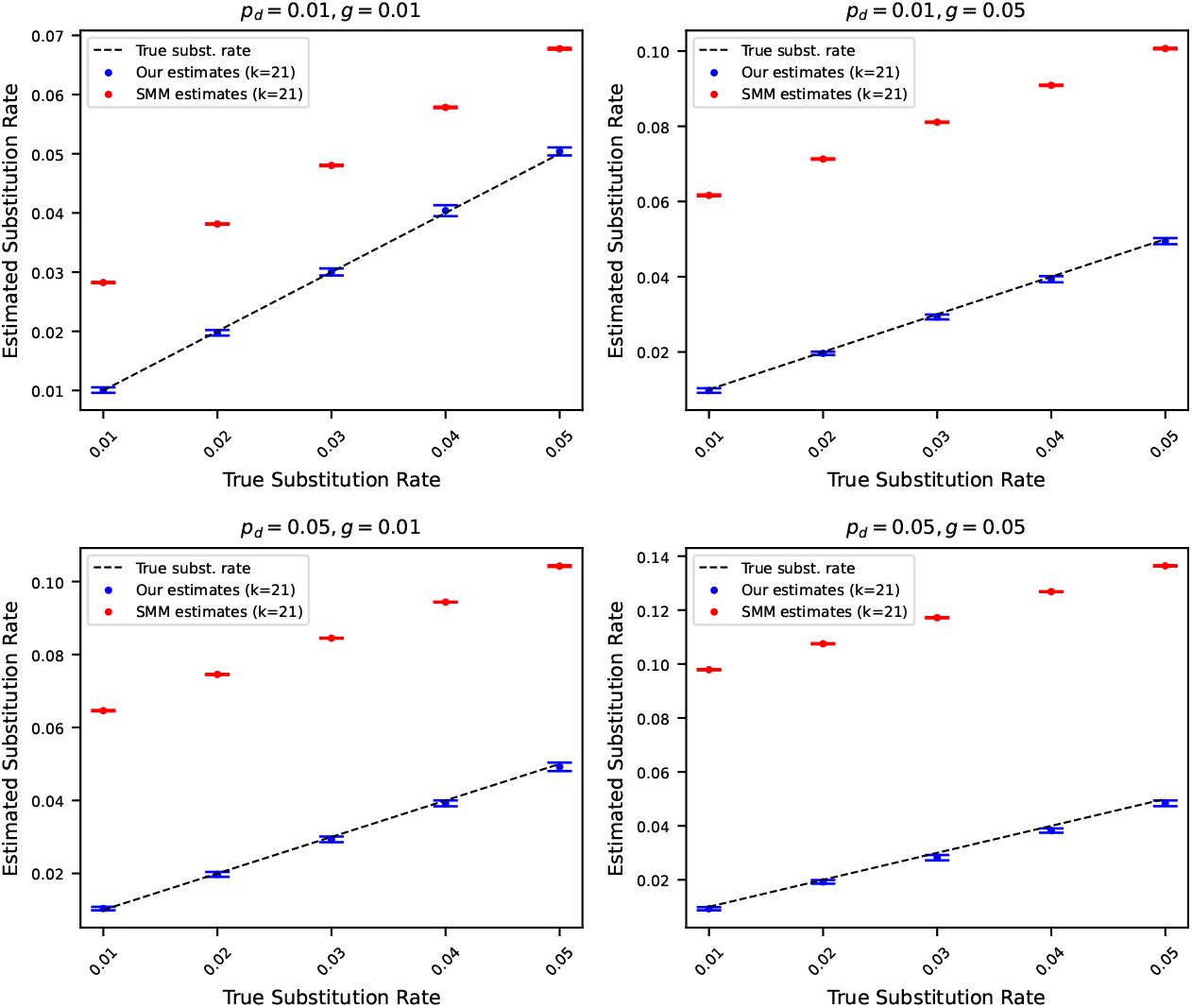
Estimated substitution rates versus true substitution rates, where the original string is the reference genome of Staphylococcus (length 2.8 million). Rates were estimated using (10) and using a simple mutation model (SMM) that only considers substitution. Each subplot corresponds to a case where *p*_*d*_ and *g* are fixed at either 0.01 or 0.05. The points show the average of 10 estimates, and the error bars show one standard deviation. The dashed black line corresponds to the true substitution rates. Estimated substitution rates using our method closely match true rates, whereas SMM overestimates due attributing to substitutions mismatching *k*-mers originating from insertions and deletions.

## 7 Conclusions

We have presented a mutation model that accommodates single-nucleotide substitutions, as well as insertions and deletions while retaining enough mathematical structure to admit closed-form rate estimators derived solely from *k*-mer statistics. From this model, we obtained algebraic estimators for the three elementary mutation rates: *p*_*s*_, *p*_*d*_, and *g* and proved relatively tight sub-exponential concentration bounds on these estimators. We also identified regimes in which the estimation becomes ill-conditioned (i.e. large *k, p*_*d*_ = 0, or sequence composition with 25% ‘A’). These results establish a bridge between sequence evolution and combinatorial word statistics, thus providing additional tools for theoretical algorithmic computational biology.

In our prototype implementation, we demonstrated that our estimates remain accurate on simulated evolution of real genomes, and outperforms a substitution-only simple mutation model by avoiding spurious attribution of indel signals. While naive counting of non-mutated and single-deletion *k*-mers sufficed to show practical accuracy of our estimators, this raises an interesting open problem: estimating the number of these non-mutated and single-deletion *k*-mers efficiently for large scale data sets.

Several directions invite further investigation. First, incorporating the count of single-substitution *k*-spans may illuminate why *p*_*s*_ remains relatively stable even for moderately large *k*. Second, our framework can extend to heterogeneous or context-dependent rates by replacing global expectations with position-specific covariates. Third, coupling our estimators with sketch-based distance measures (such as in [11]) may provide a theory-backed avenue for larger scale applications such as phylogenetic placement in the presence of high indel activity. Finally, a more thorough investigation on real genomic data (where the unitig-based approach we used in the practical implementation starts to become infeasible) will be necessary to understand the utility of the mutation estimates in practice.

In summary, by utilizing probabilistic modeling and concentration inequalities, we provide a theoretical foundation and initial practical implementation for quantifying the parameters of a relatively complex mutation process directly from *k*-mers. We anticipate that these ideas will continue to inform new alignment-free computational biology tasks, particularly relevant as sequencing data continues to outpace traditional alignment-based paradigms.

## Related Version

A preliminary version of this paper appeared in the proceedings of WABI 2025 [10].

## A Standard concentration inequalities

This section gathers several definitions and standard concentration inequalities that we use in our proofs. Certain sums of independent random variables exhibit sharp concentration, and this is captured by the following standard inequalities. The first one is the Chernoff bound for sums of independent and identically distributed Bernoulli random variables [1, 4].

### ►Lemma 8.

*Let X*_1_, …, *X*_*n*_ *be a sequence of independently and identically distributed Bernoulli random variables. Consider* 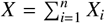 *and let* **E**[*X*] = *µ*. *Then, the following three inequalities hold:*

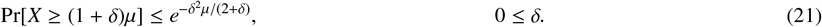

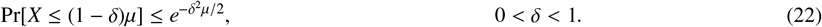

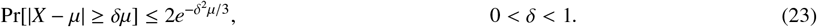

A more general inequality that applies to sums of independent and bounded random variables is Hoeffding’s inequality [12].

### ►Lemma 9.

*Let X*_1_, …, *X*_*n*_ *be a sequence of independent random variables taking values in* [*a, b*]. 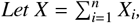 *and* **E**[*X*] = *µ*. *Then, for any δ* > 0:

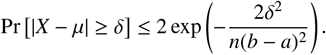

We will also analyze sums of certain *unbounded* random variables. For this, we recall the definition of sub-exponential random variables. For a comprehensive survey about this type of random variables, the reader is referred to [24, Chapter 2.7].

### Definition 10.

*Let a, b be fixed positive real numbers. A random variable X such that* **E**[*X*] = 0 *is called* (*a, b*)*-sub-exponential if for all t* ≥ 0

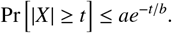

For sums of (*a, b*)-sub-exponential random variables, the following version of Bernstein’s inequality can be derived following the argument in the proof of [24, Theorem 2.8.1].

### ►Lemma 11.

*Let a, b be fixed positive real numbers and let X*_1_, …, *X*_*n*_ *be a sequence of independent* (*a, b*)*-sub-exponential random variables*.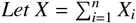. *Then, for every t* ≥ 0:

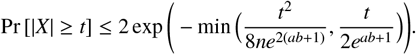

For completeness, we provide a proof of this concentration inequality in Appendix B. We derive and use this variant of Bernstein’s inequality instead of the one in [24, Theorem 2.8.1] since that one does not track the dependency on *a* and *b* in the tail bound.

We will also use the following lemma to combine concentration inequalities, with a proof given in Appendix C.

### ►Lemma 12.

*For all random variables X*_1_, …, *X*_*n*_ *and numbers t*_1_, …, *t*_*n*_,

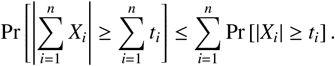

## B Proof of main result

### ►Lemma 1.

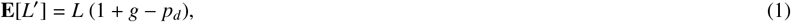

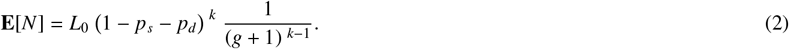

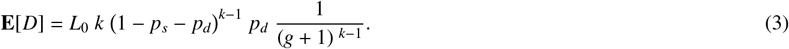

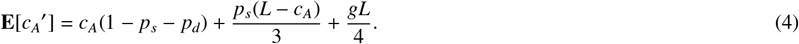

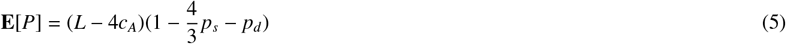

**Proof**. *Proof of* (1): Let *W*_*i*_ be an indicator random variable that is 1 in the event that character *S* _*I*_ was deleted. Then 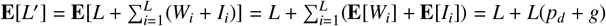.

*Proof of* (2): Let *N*_*i*_ be the indicator that *k*-span *K*_*i*_ has no mutations. This happens when each of its *k* positions has a Stay operation and there are no string inserted inside the *k*-span, i.e. *l* _*j*_ = 0 for each *j* ∈ (*i* + 1, …, *i* + *k* − 1). This happens with probability (1 − *p*_*s*_ − *p*_*d*_)^*k*^/(*g* + 1)^*k*−1^. The result follows since 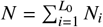.

*Proof of* (3): Let *D*_*i*_ be an indicator that *K*_*i*_ has a single deletion and no other mutation. This happens when there are no strings inserted inside the *k*-span, one position has a deletion, and *k* − 1 other positions have a Stay operation. The single deletion operation could be at any of the *k* positions. Then, **E**[*D*_*i*_] = *k*(1 − *p*_*s*_ − *p*_*d*_)^*k*−1^ *p*_*d*_/(*g* + 1)^*k*−1^. The result follows since 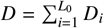.

*Proof of* (4): Let *X*_*i*_ be an indicator *S* _*i*_ is not deleted or substituted, which happens with probability 1 − *p*_*s*_ − *p*_*d*_. Let *Y*_*i*_ bet the indicator *S* _*i*_ was substituted into ‘A’, which happens with probability *p*_*s*_/3. Let *Z*_*i*_ be the number of ‘A’s in the inserted string associated with position *i*. We have that

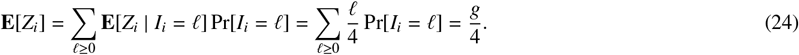

We can express *c*_*A*_′ as: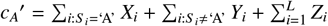, and thus get the claimed result that

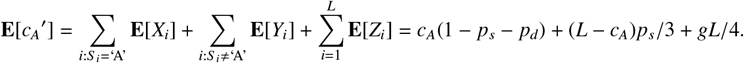

*Proof of* (5): It follows directly by linearity of expectation and (1) and (4). ◀

### ►Lemma 4.

*Suppose that* 4*c*_*A*_ < *L. For any δ* ∈ (0, 1), *all the following hold:*

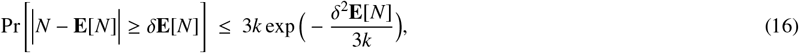

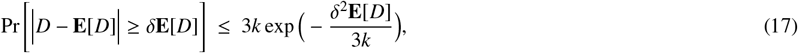

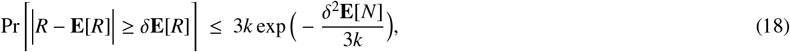

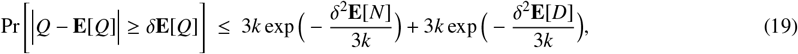

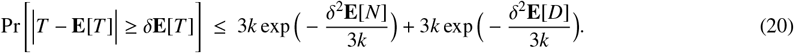

**Proof**. *Proof of* (16): Recall from the proof of Lemma 1 that we can express 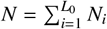 where *N*_*i s*_is the indicator that *K*_*i*_ has no mutations. The *N*_*i*_’s are not independent, but *N*_*i*_ and *N*_*j*_ are independent if |*i* − *j*| ≥ *k*. Then, for *𝓁* ∈ {1, …, *k*}, define

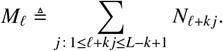

Observe that the *M*_*𝓁*_’s are defined such that each is a sum of independent identically distributed Bernoulli random variables with success probability *q* = (1 − *p*_*s*_ − *p*_*d*_)^*k*^/(*g* + 1)^*k*−1^ and 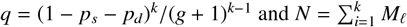. The Chernoff bound in (23) implies that for any *δ* ∈ (0, 1),

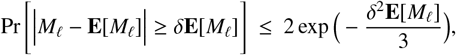

Using the fact that 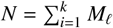 and Lemma 12, we get

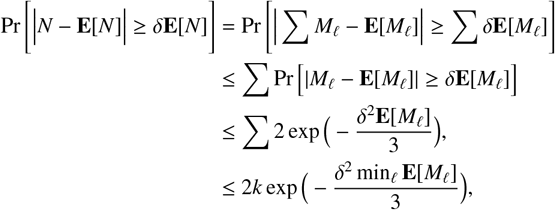

Finally, observe that each *M*_*𝓁*_ is the sum of at least 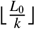 Bernoulli variables. Then 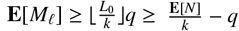. Using the fact 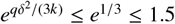, we get

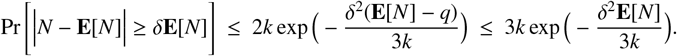

*Proof of* (17): The concentration of *D* is proven in an identical manner to the concentration of *N*. Recall from the proof of Lemma 1 that we can express 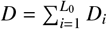, where *D*_*i*_ is the indicator that the *K*_*i*_ has no mutations except exactly one deletion. These *D*_*i*_s have the same dependence structure as the *N*_*i*_s, so the proof for the concentration of *D* is identical to the one for *N*.

*Proof of* (18): Since we assumed 4*c*_*A*_ < *L*, we can multiply both sides of the inequality inside the probability of (16) by *k*(*L* − 4*c*_*A*_) to obtain (18).

*Proof of* (19): The concentration from *Q* comes by using Lemma 12 to combine the the concentration inequalities of *N* and *D*. That is,

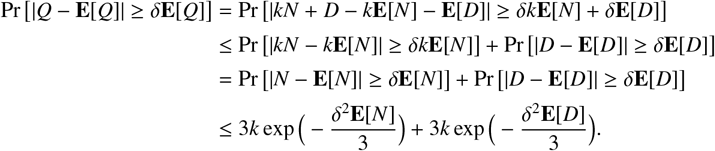

*Proof of* (20): The concentration of *T* is derived in a way identical to the concentration of *Q*. ◀

### ►Lemma 5.

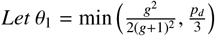. *For any δ* ∈ (0, 1):

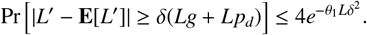

**Proof**. Let *D*_*i*_ be an indicator that is 1 in the event that the operation at *i* was Del. Let us define 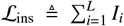, and 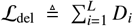, so that *L*′ = *L* + ℒ_ins_ − ℒ_del_. We will obtain the desired concentration inequality for *L*′ from separate concentration inequalities for ℒ_ins_ and ℒ_del_.

First, notice that ℒ_ins_ ≥ (1 + *δ*)*Lg* is equivalent to observing at least (1 + *δ*)*Lg* failures before observing *L* successes from a series of independent Bernoulli trials with success probability 1/(*g* + 1).

This is in turn equivalent to observing at most *L* successes in (1 + *δ*)*Lg* + *L* independent Bernoulli trials with success probability 1/(*g* + 1) each. Hence, if {*Z*_*i*_}_*i*≥1_ is a sequence of independent Bernoulli random variable with success probability 1/(*g* + 1), we have

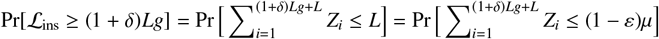

where

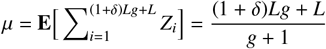

and 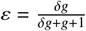. From the Chernoff bound in (22) we obtain

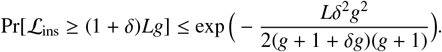

Proceeding in an analogous manner and using the Chernoff bound of (21), we can prove the probability for the other tail

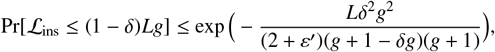

where 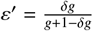. From a union bound we deduce that:

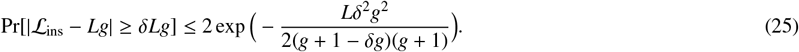

To establish the concentration of ℒ_del_, note that this random variable is a sum of *L* independent Bernoulli random variable with parameter *p*_*d*_. Then, **E**[ℒ_del_] = *Lp*_*d*_ and the Chernoff bound in (23) yields that:

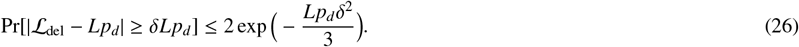

Using Lemma 12, we now combine the concentration bounds of (25) and (26) into a concentration bound for *L*′:

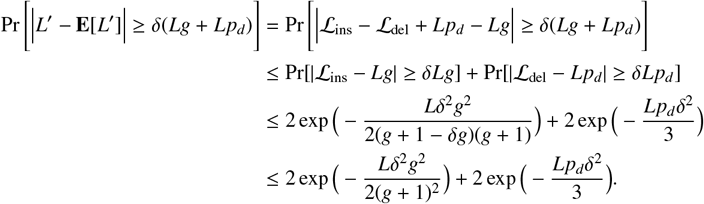

Setting 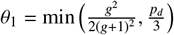, we obtain:

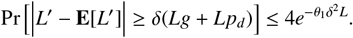

◂

### ►Lemma 6.

*Let θ*_2_ = max(*g* + 1, 8). *Then, the following holds for any δ* > 0:

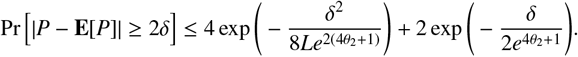

*If we also assume that* 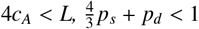, *and for any δ*_0_ ∈ (0, 1) *then*

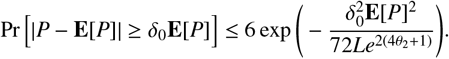

**Proof**. We will consider the contributions to *P* = *L*′ − 4*c*_*A*_′ from the characters in *S* and from newly inserted characters separately. For this, we define the random variable *X*_*i*_ that is 0 if *S* _*i*_ is deleted, it is −3 if *S* _*i*_ is not deleted and is an ‘A’ after the mutation process, and it is 1 otherwise.

We also define the random variable *Z*_*i*_ = *I*_*i*_ − 4*C*_*i*_ where recall that *I*_*i*_ is the length of the inserted string at position *i* and *C*_*i*_ is the random variable corresponding to the number of ‘A’s in the inserted string at position *i*. Note that each inserted character contributes 1 to *P*, if it is a non-’A’, or contributes−3 to *P*, if it is a ‘A’. We can now write 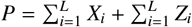. For ease of notation, let 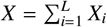 and 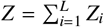.

Observe that the values that *X*_*i*_ can take on are limited to the range of [−3, 1]. Using this and the fact that the *X*_*i*_s are independent of each other, we apply Hoeffding’s inequality (Lemma 9) to get that for any *δ* > 0,

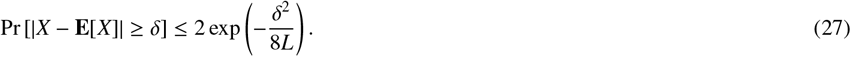

To show that *Z* is strongly concentrated around its mean, we first prove that the *Z*_*i*_’s are sub-exponential random variables and then use Bernstein’s inequality (Lemma 11).

### ►Lemma 13.

*Let θ*_2_ = max(*g* + 1, 8). *For each Z*_*i*_, **E**[*Z*_*i*_] = 0 *and, for all t* ≥ 0:

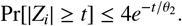

We prove this lemma shortly, but first we finish the proof of Lemma 6. As the *Z*_*i*_’s are mean-zero (4, *θ*_2_)-sub-exponential random variables, it follows from Lemma 11 that for any *δ* > 0,

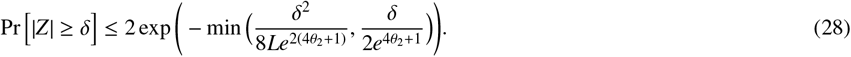

We now use Lemma 12 to combine the concentrations of *X* and *Z* into a concentration for *P*. For any *δ* > 0,

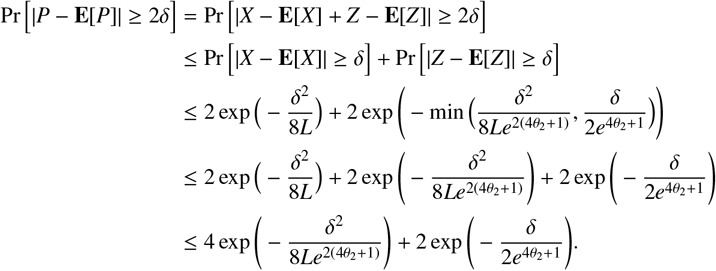

Further, applying the assumptions that 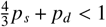 and 4*c*_*A*_ < *L* to the expected value of *P* given by (5), we have that 0 < **E**[*P*] < *L*. Therefore, given *δ*_0_ ∈ (0, 1), we can plug *δ* = *δ*_0_**E**[*P*]/2 into the above to obtain

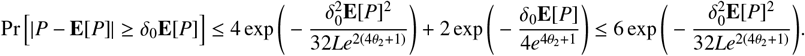

◂

**Proof of Lemma 13**. Recall that *Z*_*i*_ = *I*_*i*_ − 4*C*_*i*_, where *I*_*i*_ is a Geometric random variable with parameter *p* ≜ 1/(*g* + 1) and mean *g*, and *C*_*i*_ is a Binomial random variable with parameters *I*_*i*_ and 1/4. Observe that **E**[*Z*_*i*_] = 0, since **E**[*I*_*i*_] = *g* and, by (24), **E**[*C*_*i*_] = *g*/4. Let *B*_*y*_ be a Binomial random variable with parameters *y* and 1/4. For *t* ≥ 0, conditioning on the value of *I*_*i*_ and using Hoeffding’s inequality (Lemma 9), we obtain:

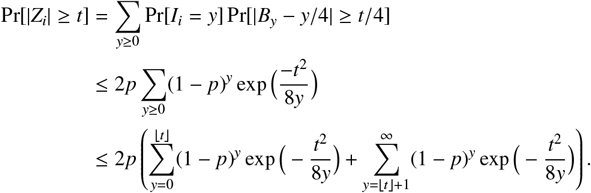

For *y* ≤ *t*, we have 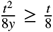, so 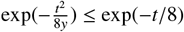 and we have

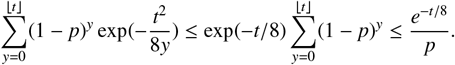

When *y* > *t*, we use the fact that *e*^−*x*^ ≤ 1 for all positive *x* and apply the formula for a geometric series; we have

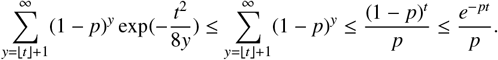

Putting these bounds together:

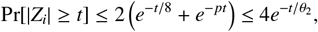

where *θ*_2_ = min{*p*, 1/8} = max(*g* + 1, 8). ◀

### ►Lemma 7.

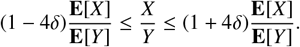

*If it also holds that* (1 − *δ*) **E**[*Z*] ≤ *Z* ≤ (1 + *δ*) **E**[*Z*], *then*

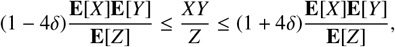

**Proof**. First, using the fact that 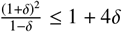, we get:

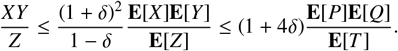

Using the fact that 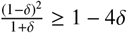, we get

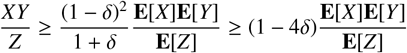

Similarly, using the facts that 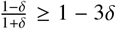 and 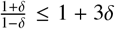, we obtain that 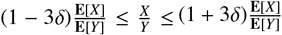.◂

### Theorem 2.

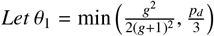*and θ*_2_= max(*g* + 1, 8). *Suppose* 4*c*_*A*_ < *L and* 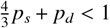. *Then, for δ* ∈ (0, 1/5):

1. 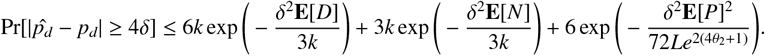
2. 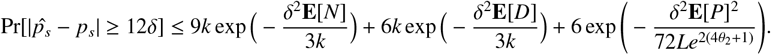
3. 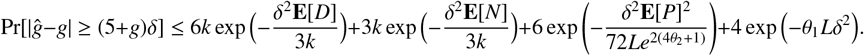

**Proof**. In Section 4, we proved the concentration bound for 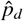 here, we will prove the bound for 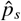 and *g*. Recall that *P* = *L*′ − 4*c*′_*A*_, *Q* = *kN* + *D, R* = *k*(*L* − 4*c*_*A*_)*N*, and *T* = 4*kN* + *D*. Let us assume our variables are indeed close to their means, i.e. for all *X* ∈ {*P, Q, R, T* }, (1 − *δ*) **E**[*X*] ≤ *X* ≤ (1 + *δ*) **E**[*X*] for *δ* ∈ (1, 1/5). Because we assume that 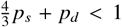 and 4*c*_*A*_ < *L*, we use (5) to obtain that **E**[*P*] > 0; in addition, Lemma 1 gives **E**[*Q*], **E**[*R*], **E**[*T*] > 0. Therefore, we can apply Lemma 7 to get

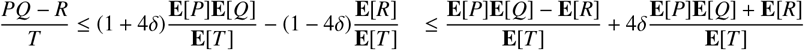

and, similarly,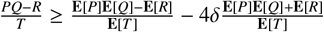. Multiplying by 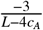 and using the definition of (13), we obtain that 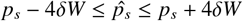 with 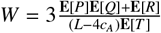. Plugging in the values given by Lemma 1, we obtain that *W* ≤ 3. Hence, 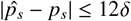. As in the proof for 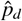, the result follows by a union bound using the individual concentration bounds for *P* (Lemma 6), *Q, R*, and *T* (Lemma 4).

To obtain the bound for *ĝ*, recall that 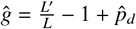. From Lemma 5, for any *δ* ∈ (0, 1)

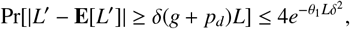

where 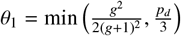. Then,

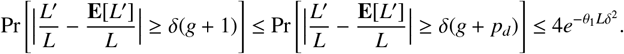

Combining this with Lemma 12, we get

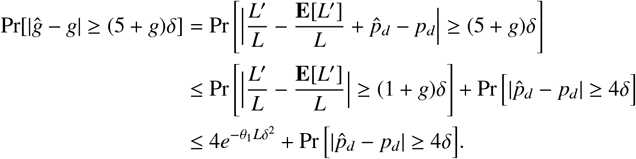

Plugging in the bounds on 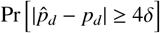 from part 2 of this theorem gives the result. ◀

## C Proofs of helper lemmas

In this section, we provide for completeness some of the proofs from Appendix A. To prove Lemma 11, we will first need the following fact.

### ►Lemma 14.

*Let a, b be fixed positive real numbers. If X is* (*a, b*)*-sub-exponential, for λ* ∈ ℝ *such that* |*λe*^*ab+1*^ | < 1/2 *we have* 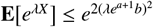.

**Proof**. Using the Taylor expansion for the exponential function, the fact that **E**[*X*] = 0, and standard inequality 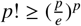, we deduce that

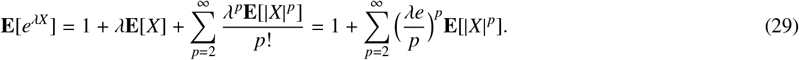

Observe that for integer *p* ≥ 1, we have the identity 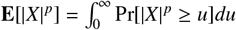, and with the change of variable *u* = *bt*^*p*^, we obtain:

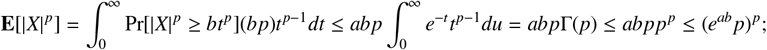

Г denotes the Gamma function and we use the fact that Γ(*x*) ≤ *x*^*x*^ for all *x* ≥ 1. Plugging this bound into (29), we obtain:

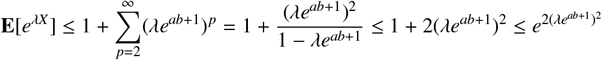

provided |*λe*^*ab*+1^| < 1/2. ◀

**Proof of Lemma 11**. Let *λ* > 0. Markov’s inequality and the independence of the *X*_*i*_s imply that:

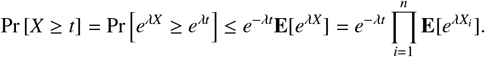

If *λ* is such that |*λe*^*ab*+1^| < 1/2, Lemma 14 implies that 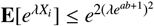 and thus

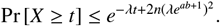

For ease of notation set *y* = *e*^*ab*+1^, and let us optimize for *λ*. The minimum of the parabola *f* (*λ*) = −*λt* + 2*y*^2^*nλ*^2^ in [0, 1/(2*y*)] aligns with is global minimum at 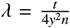 when 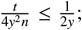 the minimum value of *f* is 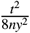 in this case. Otherwise, if 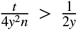, the the minimum occurs at the boundary point 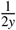 and has value 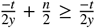. Combining these two bounds:

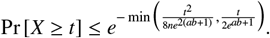

Repeating the same argument, we obtain the same bound for Pr[−*X* ≥ *t*] and the result follows from a union bound. ◀

**Proof of Lemma 12**. First observe that |Σ*X*_*i*_ | ≤ Σ|*X*_*i*_ | and therefore |Σ*X*_*i*_ | ≥ Σ*t*_*i*_ implies that Σ|*X*_*i*_ | ≥ Σ*t*_*i*_.Hence Pr[|*X*_*i*_| ≥ Σ*t*_*i*_] Pr[Σ|*X*_*i*_ | ≥ Σ*t*_*i*_]. Second, Σ|*X*_*i*_ | ≥ Σ*t*_*i*_ implies that there is at least one *i* such that |*X*_*i*_| ≥*t*_*i*_. Therefore, Pr[Σ|*X*_*i*_ | ≥ Σ*t*_*i*_] ≤ Pr[|*X*_1_| ≥*t*_1_⋃…⋃ |*X*_*n*_| ≥*t*_*n*_] Third,by the union bound, Pr[|*X*_1_| ≥*t*_1_⋃…⋃ |*X*_*n*_| ≥*t*_*n*_] ≤ Σ Pr[|*X*_*i*_ | ≥*t*_*i*_]. Combining the three observations in sequences, we get the result. ◀

